# CRIPTO promotes extracellular vesicle uptake and activation of cancer associated fibroblasts

**DOI:** 10.1101/2024.03.01.583059

**Authors:** David W. Freeman, Brooke L. Gates, Mauri D. Spendlove, H. Evin Gulbahce, Benjamin T. Spike

## Abstract

Expression of CRIPTO, a factor involved in embryonic stem cells, fetal development, and wound healing, is tied to poor prognosis in multiple cancers. Prior studies in triple negative breast cancer (TNBC) models showed CRIPTO blockade inhibits tumor growth and dissemination. Here, we uncover a previously unidentified role for CRIPTO in orchestrating tumor-derived extracellular vesicle (TEV) uptake and fibroblast activation through discrete mechanisms. We found a novel mechanism by which CRIPTO drives aggressive TNBC phenotypes, involving CRIPTO-laden TEVs that program stromal fibroblasts, toward cancer associated fibroblast cell states, which in turn prompt tumor cell invasion. CRIPTO-bearing TEVs exhibited markedly elevated uptake in target fibroblasts and activated SMAD2/3 through NODAL-independent and - dependent mechanisms, respectively. Engineered expression of CRIPTO on EVs enhanced the delivery of bioactive molecules. *In vivo*, CRIPTO levels dictated TEV uptake in mouse lungs, a site of EV-regulated premetastatic niches important for breast cancer dissemination. These discoveries reveal a novel role for CRIPTO in coordinating heterotypic cellular crosstalk which offers novel insights into breast cancer progression, delivery of therapeutic molecules, and new, potentially targetable mechanisms of heterotypic cellular communication between tumor cells and the TME.

## MAIN TEXT

Tumors are comprised of an intricate network of parenchymal and stromal cells, whose intercellular signaling governs cell state change, resistance to therapy, and tumor growth at both primary and distant sites^1^. Deciphering the molecular underpinnings of coordinated heterotypic signaling in multi-cellular tumor microenvironments (TME) is crucial for understanding tumor progression, and could reveal effective therapeutic strategies aimed at non-autonomous drivers of cancer progression and metastasis.

CRIPTO is a small GPI-anchored signaling protein encoded by the *TDGF1* gene that balances TGF-β singaling and independent pathways that activate c-Src/MEK/AKT^2–4^. It is prominently expressed in stem cells and early development, largely undetectable in adult homeostatic tissue, but reexpressed in the context of oncogenesis and wound healing^4^. In mammary developmental studies, inhibition of CRIPTO with a soluble CRIPTO ligand trap, Alk4^L75A^-Fc (A4Fc), significantly impaired the multipotency of murine mammary stem cells and their proliferative renewal capacity^5^. Elevated CRIPTO is detectable in nearly 80% of breast cancers and its overexpression in upwards of 50% of invasive breast tumors is associated with poor prognosis and advanced disease states^6,7^. Knocking down CRIPTO, blocking its signaling with A4Fc, or delivering toxic payloads to CRIPTO expressing cells with an antibody-drug conjugate reduces primary tumor growth and distant metastasis in triple negative breast cancer (TNBC) xenograft models^8,9^.

Though CRIPTO is known to act both autonomously *and* as a soluble secreted factor, it has been studied in cancer primarily as an autonomous factor within the epithelial tumor parenchyma. However, there is good reason to suspect CRIPTO may coordinate the activity of multiple cooperating cell types, given its correlation to injury related fibrosis in experimental models and our prior identification of CRIPTO-dependent effects on mammary stem cells cocultured with fibroblasts^10,5^. Coordination of epithelial fibroblast crosstalk is also critical in cancer. Breast cancer-associated fibroblasts (CAFs) are known to facilitate treatment resistance, tumor proliferation, and enable metastatic outgrowth, partly through mechanical alterations of the ECM and cytokine secretion, though key targetable regulators remain to be identified^1,11–13^.

Here, we demonstrate that CRIPTO governs a novel communication axis between tumor cells and fibroblasts involving extracellular vesicles (EVs). This discovery emerged from our analysis of single cell profiles of the TNBC cell line, MDA-MB-468. Specific subpopulations of MDA-MB-468 cells grown as organoids were found to be CRIPTO-dependent and to bear gene expression signatures, distinct from classic CAF or EMT, that correlate with CAFs in archival patient data. Rather, their enrichment for vesicular trafficking and extracellular vesicle gene programs prompted us to evaluate novel mechanisms of CRIPTO-mediated tumor cell-fibroblast crosstalk and the potential role of EVs.

EVs are secreted mediators of intercellular crosstalk capable of shuttling proteins, DNA/RNA, lipids, and various metabolites. Classically, EVs have been divided into three classes: (1) exosomes formed through endocytic intermediates, (2) microvesicles produced through outward membrane blebbing, and (3) apoptotic bodies generated from dying cells^14^. Roles for EVs in tumor initiation and progression have been described, including their capacity to prime premetastatic niches in distant organs for tumor cell colonization^15^. We demonstrate that tumor cell derived EVs (TEVs) bearing CRIPTO act as paracrine fibroblast programming factors.

Mechanistically, CRIPTO-laden EVs can activate NODAL signaling in target fibroblasts while exhibiting enhanced NODAL-independent EV uptake *in vitro,* and *in vivo* in lungs of mice where EVs have been implicated in fibroblast programming to form pre-metastatic niches^15^. This discovery represents a key milestone in understanding CRIPTO activity in multicellular contexts, including tumor microenvironments where CRIPTO-dependent EV signaling could represent a novel therapeutic target for breast cancer and beyond.

### Tumor cell-derived CRIPTO activity correlates with CAFs

We previously showed that CR1 antagonism by A4Fc limits various phenotypes in TNBC cells associated with stemness and tumor progression, including the capacity of TNBC (MDA-MB-468) organoids to undergo serial passaging despite no overt pre-passage organoid phenotypes^9^. This suggested CRIPTO may regulate the maintenance of a minority stem-like cell population required for secondary organoid outgrowth^9^. To test this, we performed single cell RNA-sequencing (scRNA-seq) of MDA-MB-468 organoids transduced with an empty tet-inducible lentiviral expression cassette (s14^Con^) or a construct with inducible expression of secreted A4Fc (s14^A4Fc^). Inhibition of CRIPTO with A4Fc induced a marked skewing in two specific cell subpopulations (denoted clusters 5 and 8) (**Fig 1A)**. s14^A4Fc^ cultures showed a marked loss of cluster 8 cells and an increase in cluster 5 cells relative to s14^Con^ cultures. We identified differentially expressed genes (DEGs) representing these populations and confirmed CRIPTO dependency of select targets using *TDGF1* knockdown (shCr) (**Fig 1B,C)**.

**Figure 1.**
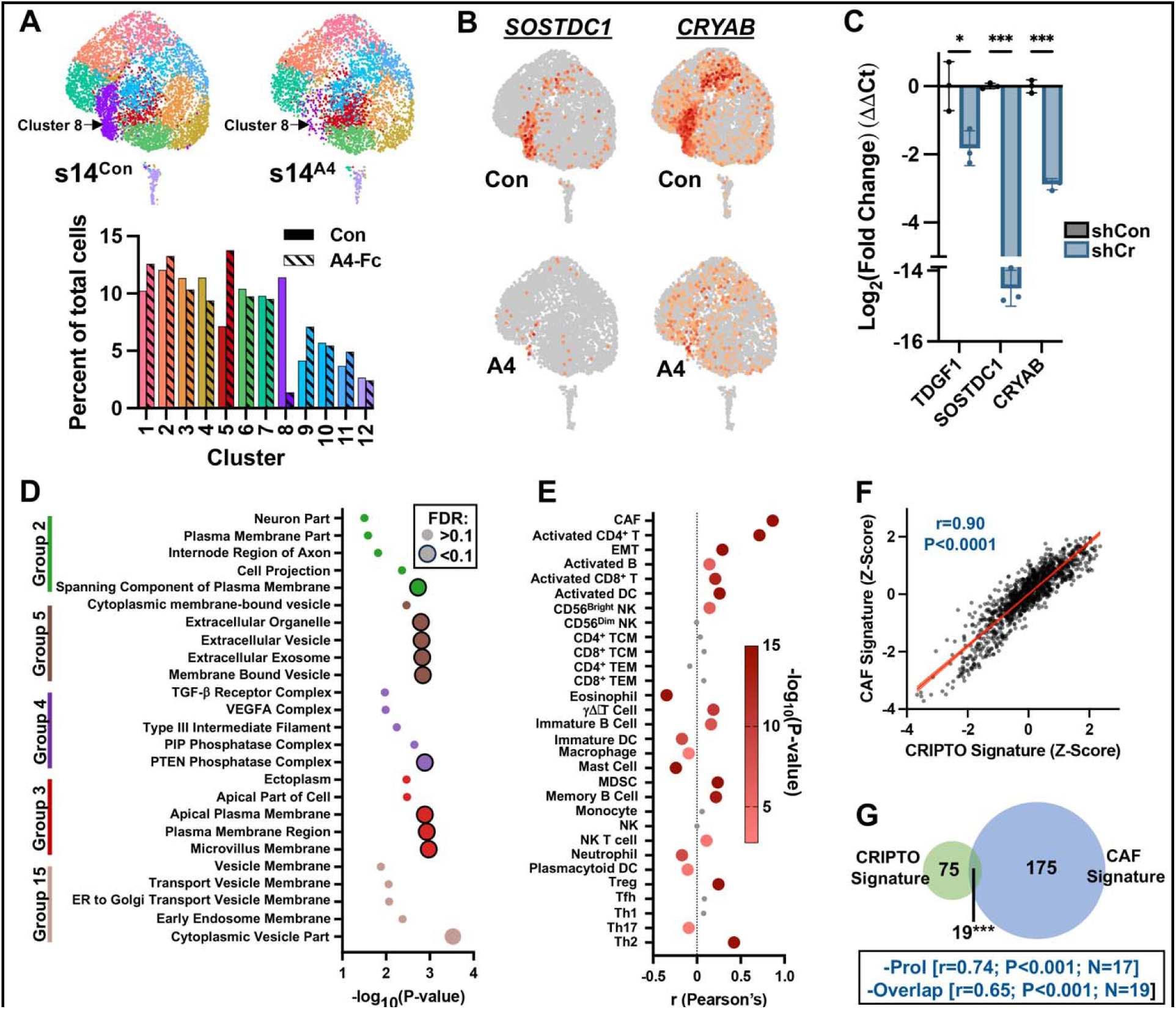
CRIPTO regulates TNBC tumor cell populations with extracellular vesicle and CAF-associated profiles. (A) UMAPS and relative cluster proportions of scRNA-seq profiles from MDA-MB-468 s14^Con^ and s14^A4Fc^ organoids. (B) Relative expression of representative cluster 8 genes. (C) RT-qPCR validation of CRIPTO-dependency of genes from B in MDA-MB-468 shCon and shCr 2D cultures; *p<0.05, ***p<0.001 Holm- Šidák Multiple T-tests. (D) Jensen cell compartment analysis of cluster 8 DEGs. Common compartments are clustered together, top 5 compartments for each enriched cluster are shown. Larger bubbles represent significantly enriched compartments (q<0.1; Benjamini-Hochberg adjusted). (E) Pearson’s correlation of CRIPTO signature with stromal cell type gene signatures, larger bubbles represent significant correlations (Bonferonni adjusted P<0.0017) (F) Pearson’s correlation between CRIPTO signature and CAF signature (TCGA Breast Cancers). (G) Overlap of genes between CAF and CRIPTO gene signatures (Hypergeometric distribution). ***p<0.001 Box=Pearson correlations and P-values when genes in common or annotated as ‘proliferation’ or ‘cell division’ by GO analysis were removed from the CRIPTO signature.

To help identify underlying biology regulated by CRIPTO in this setting, we examined the enrichment of DEGs associated with these subpopulations in curated gene sets for biological processes and cellular compartmentalization^16,17^. Genes characteristic of cluster 8 showed enrichment in factors predicted to localize to extracellular vesicles (EV) and exosomes **(Fig 1D, Fig S1A,B).** Though somewhat unexpected based on CRIPTO’s role as a cell surface coreceptor, this link between CRIPTO and EV biology was consistent with punctate staining we previously observed for CRIPTO in tumor and developmental settings^5,9^, unpublished proteomics work from our group analyzing A4Fc pulldowns from xenografts indicating associated EV-markers, and recent reports of CRIPTO expression in EVs from cholangiocarcinoma patients^18^.

Given the role of EVs as critical signaling mediators between tumor cells and tumor-supportive stroma, we investigated whether EVs drive ‘call and response’ programs between CRIPTO-regulated tumor cell populations and TME cell types^15,19^. First, we generated a composite gene signature (CRIPTO Signature) for cluster 8 and cluster 5 reflecting tumor cell states under CRIPTO’s influence and assessed its correlation with TME cell populations using TCGA data (**Fig 1E, Table S1**)^20,21^. The CRIPTO signature was overexpressed in TNBC relative to other subtypes, and among stromal signatures in the TCGA database, was most strongly correlated with a CAF gene signature, an association that persisted even after adjusting for the modest number of shared genes between the signatures and proliferation-related genes (**Fig 1E-G; Fig S1C-E)**. These findings are consistent with a CRIPTO-regulated tumor subpopulation influencing stromal fibroblast programs within the TME.

### CRIPTO drives EV uptake by recipient fibroblasts

Recent studies have identified CRIPTO in TEVs in several cancer types^18,22^. To determine whether CRIPTO was likely to be present in EVs within the TME of breast cancers in our system, we reanalyzed MDA-MB-468 xenograft tumors for expression and localization of CRIPTO and the canonical EV marker CD81^23^. CRIPTO colocalized with CD81 in discrete punctate structures along hypoxic tumor borders of MDA-MB-468 xenografts, where we previously observed CRIPTO-dependent tumor cell phenotypes (**Fig 2A**)^9^.

**Figure 2.**
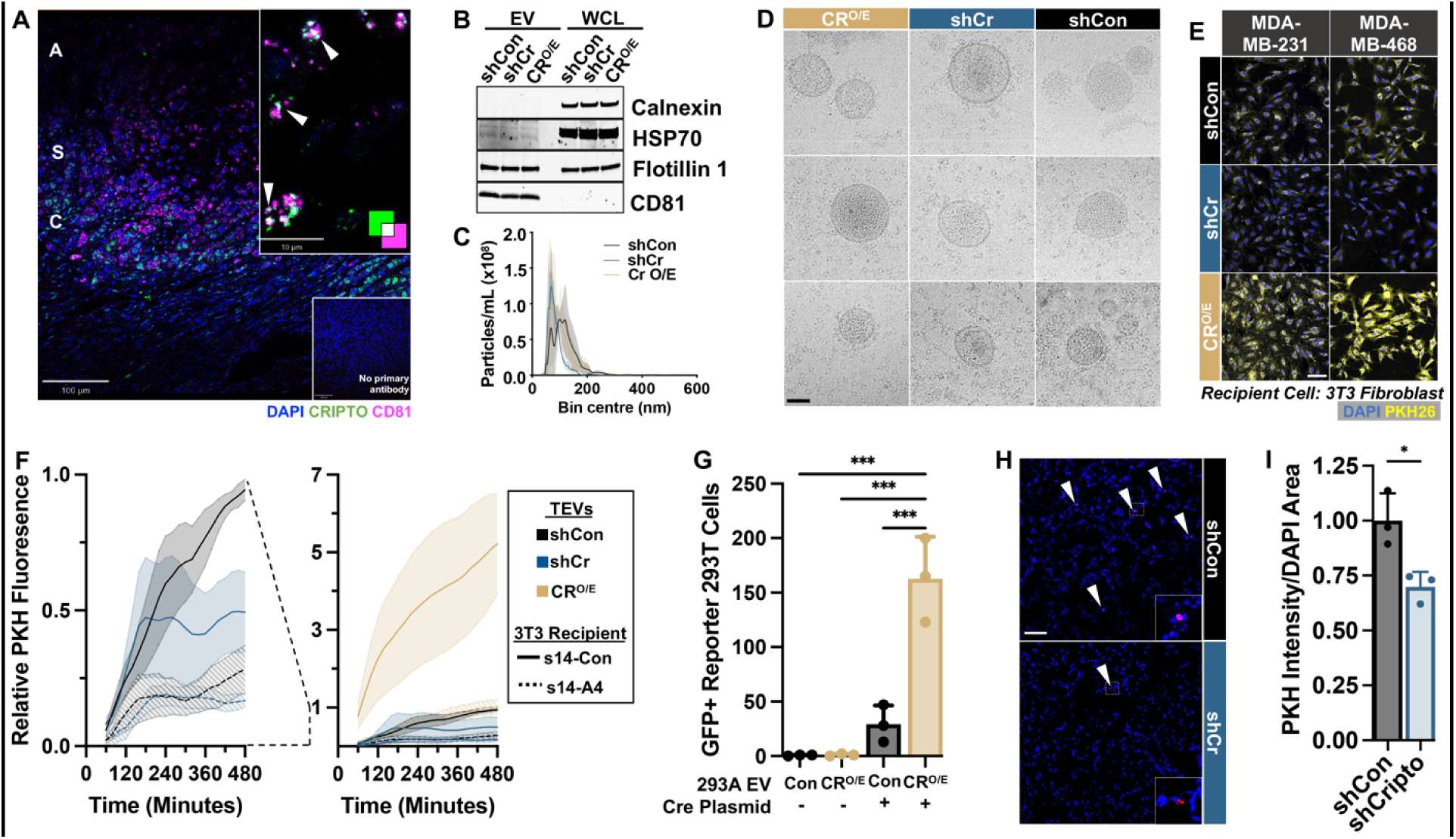
CRIPTO promotes EV uptake *in vitro* and *in vivo*. (A) Immunofluorescent co-localization of CD81 and CRIPTO in MDA-MB-468 xenograft tumor; A= acellular, S= stressed, and C= cellular regions. (B) Purity of TEVs isolated from MDA-MB-468 shCon, shCr and Cr O/E by Western blot. (C) Nanoparticle tracking analysis of 0.2μm filtered EV. (D) Representative cryo-electron micrographs of EVs from different CRIPTO-modulated cells, scale bar = 50nm. (E) Representative images of PKH labeled TEVs from MDA-MB-468 and -231 shCon, shCr, and Cr O/E cells. Scale Bar=100μm. (F) Moving average fluorescent intensity of 3T3^s14-Con^ or 3T3^s14-A4^ cells treated with PKH26 labeled TEVs from MDA-MB-468 shCon, shCr, and Cr O/E cells. (G) Number of GFP+ floxed red-to-green reporter 293T cells after treatment with CRE-carrying EVs from Con or CR^O/E^ 293A cells. N=3; One-way ANOVA with Tukey’s multiple comparisons, ***P<0.001. (H) Representative images and (I) Quantification of PKH intensity/cell in lungs from mice treated with either PKH26 labeled EVs isolated from shCon or shCr cells (N=3 animals/group; 5 field of views/animal). Two-tailed student’s T-test, *P<0.05.

Indeed, isolated EVs from conditioned media of shCon MDA-MB-468 *in vitro* cultures contained CRIPTO protein but otherwise exhibited no obvious difference in canonical EV markers, size distribution, production quantity, or morphology from EVs isolated from shCr or CRIPTO overexpressing (CR^O/E^) cultures (**Fig S1F, Fig 2B-D**). Nevertheless, microscopy of fibroblasts treated with TEVs produced from shCon, shCr, or CR O/E MDA-MB-468 and -231 cells demonstrated a marked difference in EV uptake (**Fig 2E**). shCr EVs were internalized at the lowest rate, while overexpression of CRIPTO dramatically increased EV internalization compared to both shCon and shCr groups in recipient fibroblast cells. Additionally, expression of A4Fc in recipient fibroblasts reduced the uptake of both shCon and CR^O/E^ TEVs from MDA-MB-468 cells, as assessed by timelapse imaging (**Fig 2F**).

Next we asked whether CRIPTO overexpression could be leveraged to promote delivery of selected payloads to recipient cells. We electroporated a Cre-expressing plasmid into EVs from the non-cancerous cell line HEK293A in which we overexpressed CRIPTO. CR^O/E^ cells enhanced deliver of the Cre-expression plasmids to recipient HEK293T cells compared to control EVs (**Fig 2G**). This highlights a therapeutic opportunity to leverage CRIPTO-mediated uptake to deliver bioactive molecules.

We then sought to determine whether CRIPTO regulates EV uptake *in vivo* in mouse lungs where premetastatic niches governed by EVs have been described and where we previously saw a drastic reduction in metastasis of the TNBC cell line MDA-MB-231 when CRIPTO was blocked by A4Fc^9,15^. Thus, we performed retroorbital injections of PKH26-labeled EVs isolated from shCon and shCr MDA-MB-231 cells and analyzed lung tissue 24 hours later. We observed EV uptake by rare cells throughout the lung in each case but noted that EVs from shCr cells exhibited significantly lower levels of uptake in the lungs than EVs from shCon cells (**Fig 2H,I**). Collectively, these data point to a previously unidentified role for CRIPTO in EV uptake.

### Paracrine CRIPTO signaling promotes reciprocal fibroblast:tumor cell crosstalk

To understand the effects of paracrine CRIPTO signaling via EVs on fibroblast activity, we first reexamined TNBC xenograft tumors expressing s14^Con^ of s14^A4Fc^ in which we had originally reported effects of A4Fc on tumor cell aggressiveness and disease progression, focusing now on the TME rather than the tumor parenchyma^9^. Consistent with a role for CRIPTO in regulating CAF activity, we noted alterations in collagen deposition under CRIPTO inhibition by A4Fc and qualitative changes in stromal cell morphology and expression of αSMA (**Fig. S2A-B**). Similarly, PDX models showing greater enrichment for the CRIPTO signature also scored higher for overall fibrosis (**Fig 3A,B**), collectively suggesting a functional relationship between CRIPTO signaling and CAF activity.

**Figure 3.**
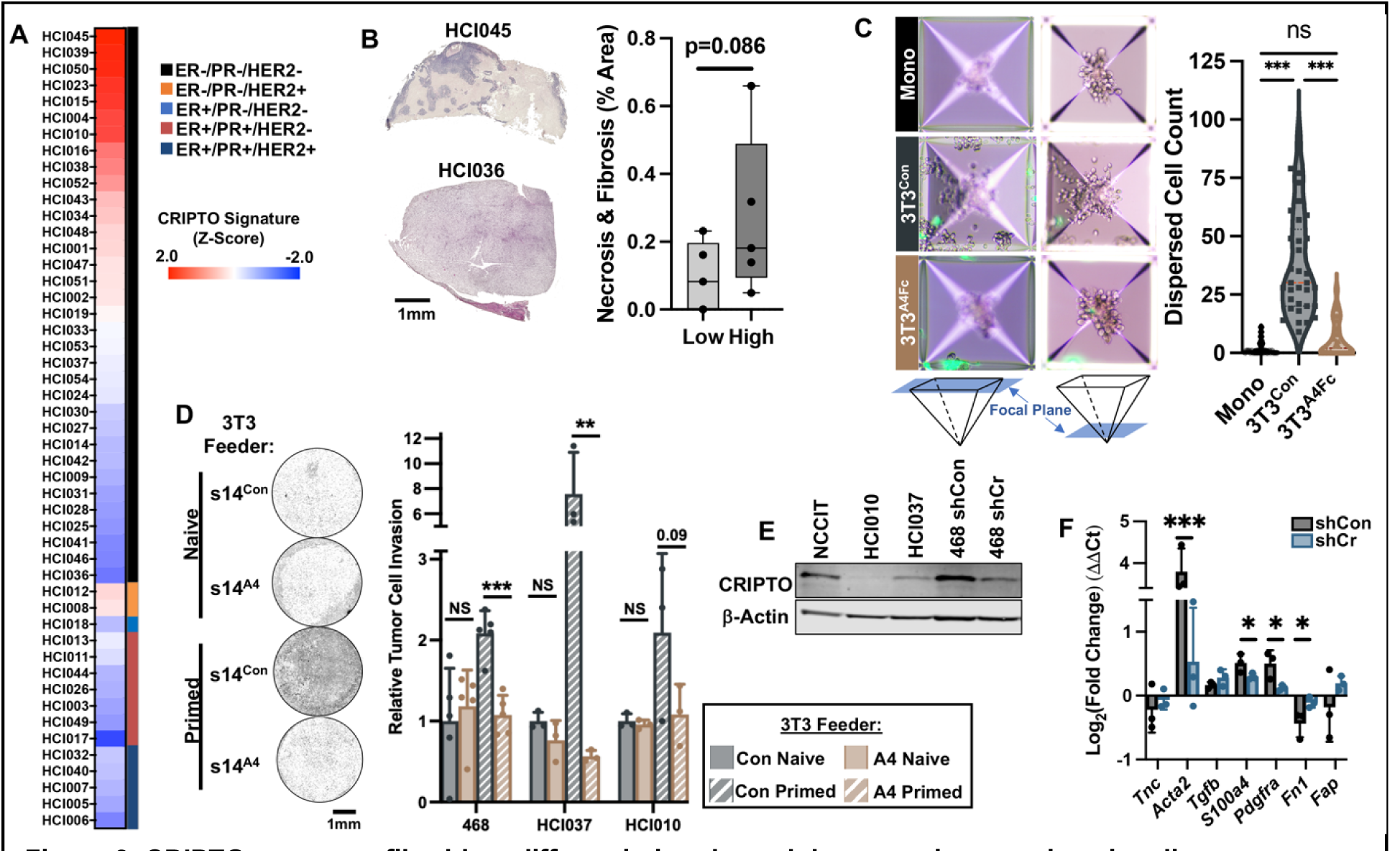
CRIPTO programs fibroblast differentiation through heterotypic paracrine signaling. (A) Relative CRIPTO signature across breast cancer PDXs. (B) Representative CRIPTO signature-high and – low PDX sections, with quantification of necrotic and fibrotic area; N=5, Student’s T-Test. Representative photos and quantification of (C) MDA-MB-468 spheroids grown with or without 3T3^s14-Con^ or 3T3^s14-A4^ fibroblasts (GFP+ cells) in microwells (N=3, 10microwells/replicate; One-way ANOVA with Tukey’s multiple comparisons) and (D) transwell invasion assay of MDA-MB-468, HCI010, and HCI037 TNBC cells grown above 3T3^s14-Con^ or 3T3^s14-A4^ fibroblasts; two-way ANOVA Šidák Multiple Comparison. **P<0.01, ***P<0.001 (E) Western blot of CRIPTO expression across cell models. (F) RT-qPCR of CAF marker genes in 3T3 fibroblasts treated with conditioned media from shCon or shCr MDA-MB-468 cells. Fold change relative to 3T3 cells treated with autologous conditioned media; Student T-test. *P<0.05 , ***P<0.001

To functionally interrogate CRIPTO-dependent coordination of tumor cell and fibroblast interaction in a controlled setting, we generated heterotypic organoid co-cultures of MDA-MB-

468 cells and growth-arrested 3T3 fibroblasts expressing s14^Con^ or s14^A4Fc^. In microwell cultures, incorporation of s14^Con^-fibroblasts stimulated tumor cell and fibroblast co-migration out of the well nadir, whereas A4Fc expression reduced tumor cell migration despite no appreciable difference in fibroblast movement (**Fig 3C**). Related experiments using transwell invasion assays confirmed that the migratory phenotype in tumor cells was at least partly driven by soluble paracrine signaling and involved tumor cell-dependent reprogramming of fibroblast cell state for reciprocal promotion of tumor cell migration. That is, when naïve tumor cells were cultured above either naïve s14^Con^ or s14^A4Fc^ fibroblasts over 48hrs, they migrated through Matrigel and the boundary membrane at low and comparable rates. However, primed fibroblasts (i.e., those previously exposed to tumor cells for a period of 3 days) dramatically enhanced the invasion of naïve tumor cells over 48 hours, and this was significantly reduced by A4Fc (**Fig 3D**). This migratory effect extended to organoid cultures of Patient Derived Xenografts (PDxOs) that endogenously expressed CRIPTO at different levels (**Fig 3D-E, Fig S2C**).

We also examined canonical CAF markers by RT-qPCR following treatment of 3T3 fibroblasts with tumor cell conditioned media (CM). Repeated treatment with CM for three days from shCon cells altered the expression of CAF markers relative to fibroblasts treated with autologous conditioned media, whereas these changes were attenuated when treating with CRIPTO knockdown CM (**Fig 3F**). Together, these data indicate tumor cell CRIPTO is indispensable in directing fibroblast reprogramming through soluble paracrine signaling.

### TEVs facilitate fibroblast activation through intercellular CRIPTO:NODAL signaling

We next examined TEVs as possible mediators of paracrine CRIPTO signaling and fibroblast programming. SMADs play a key role in CAF activation and are regulated by various TGF-β family ligands and CRIPTO^24,25^. We, therefore, investigated SMAD2/3 activation in fibroblasts as a function of paracrine signaling by CRIPTO-bearing TEVs. Conditioned media from control MDA-MB-468 cells potently induced SMAD2/3 phosphorylation in fibroblasts, and the activity was specifically associated with the isolated EV fraction of the donor cell CM from controls, while it was absent from EV depleted fractions and from all fractions of CM isolated from shCr cells (**Fig 4A**). This implicates TEVs as the predominant signaling mechanism mediating non-autonomous CRIPTO signaling between tumor cells and fibroblasts. RNA sequencing of fibroblasts treated with EVs from shCon, shCr or CRIPTO overexpressing MDA-MB-468 cells revealed CRIPTO overexpression upregulated SMAD target genes, as well as genes enriched in the matrisome, ER stress response pathways, and distinct fibroblast cell-types in archival datasets (**Fig S2D-E**).

**Figure 4.**
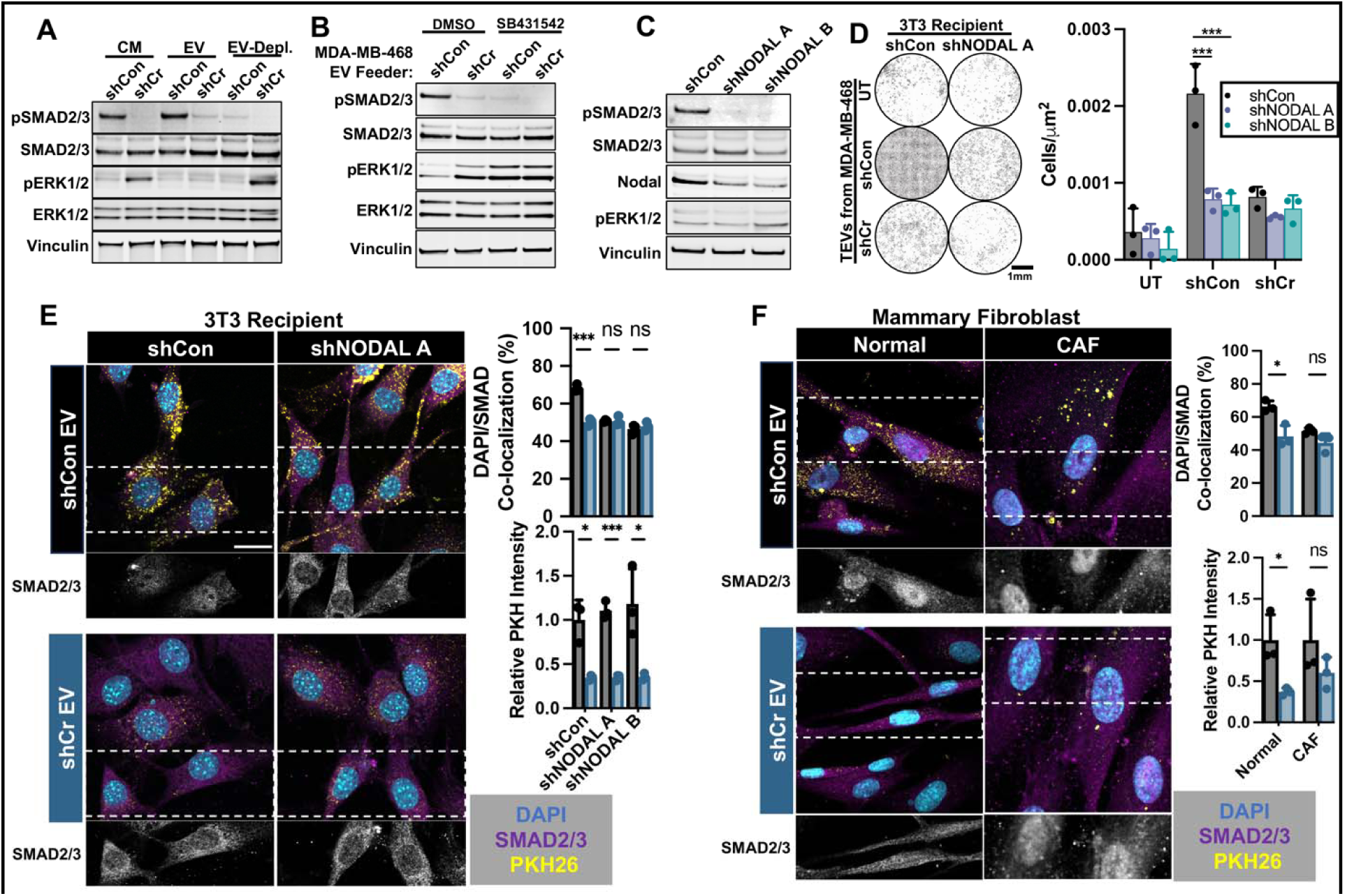
CRIPTO promotes SMAD activation and EV uptake in fibroblasts through discrete NODAL-dependent and –independent pathways. Western blots of 3T3s treated with (A) conditioned media (CM), TEVs or TEV-depleted media (EV-depl) collected from shCon and shCr MDA-MB-468 cells and (B) TEVs from MDA-MB-468 shCon or shCr cells in addition to treatment with ALK4,5,7 inhibitor, SB431542 (10μM, 16hrs). (C) Western blot of control (shCon) and NODAL knockdown (shNODAL) in 3T3s treated with TEVs from MDA-MB-468 shCon cells. (D) Invasion assay of MDA-MB-468 cells cultured above shCon or shNODAL 3T3s treated with EVs from MDA-MB-468 shCon or shCr cells. (E) Immunocytochemistry of shCon and shNodal 3T3s and (F) primary mammary fibroblasts and CAFs treated with PKH26 labeled EVs isolated from MDA-MB-468 shCon or shCr cells. Quantification of nuclear SMAD2/3 and relative PKH intensity on the right (N=3; 4 fields of view/replicate); Holm-Šidák Multiple T-tests. Scale Bar = 20μm. *P<0.05 , **P<0.01, ***P<0.001

CRIPTO acts through ALK4 in conjunction with various TGF-β family ligands to modulate SMAD2/3 activation^26^. We confirmed the invovment of ALK4,5,7 receptor complexes in CRIPTO mediated SMAD2/3 activation in fibroblasts using pharmacological inhibition (SB431542). ALK4,5,7 inhibition in fibroblasts blocked pSMAD2/3 increases following treatment with shCon TEVs (**Fig 4B**).

CRIPTO-mediated activation of SMAD2/3 typically involves the TGF-β family ligand NODAL and NODAL has previously been associated with active CAFs in TNBC^27^. We asked whether CRIPTO-expressing TEVs promoted SMAD2/3 activation by facilitating signaling through locally produced NODAL in fibroblasts. We found NODAL knockdown (shNODAL) in recipient fibroblasts blocked SMAD2/3 phosphorylation in fibroblasts treated with TEVs from CRIPTO expressing TNBC cells (**Fig 4C**). These data suggested a model in which tumor cells provide CRIPTO to fibroblasts in the form of CR-laden EVs that then act in concert with fibroblast-expressed NODAL to program fibroblasts for reciprocally promotion of tumor cell invasion. Consistent with this conjecture, fibroblasts treated with isolated shCon TEVs promoted tumor cell invasion and this effect was reversed by knockdown of CRIPTO in the TEV-producing tumor cells or by NODAL knockdown in the recipient fibroblasts (**Fig 4D**). These data highlight an intercellular signaling axis dependent on CRIPTO and NODAL expression in separate cell populations.

We next asked whether TEV uptake was contingent on NODAL expression within fibroblasts. We found TEV uptake was NODAL-independent as fluorescently labeled TEVs from shCon cells were taken up equally by control and shNODAL fibroblast while TEVs from shCr cells were internalized less readily than controls, regardless of NODAL expression in fibroblasts. In contrast, and consistent with western analysis, levels of nuclear SMAD2/3 (indicative of SMAD activation) were dependent on both fibroblast NODAL levels and tumor CRIPTO levels (**Fig 4E**). Critically, the differential EV uptake observed in our 3T3 model system was recapitulated using primary human mammary fibroblasts as the recipient cell type. That is, in primary human fibroblasts, shCon EVs were taken up more readily than shCr EVs and promoted nuclear localization of SMAD2/3 that was diminished when using shCr-donor cell EVs (**Fig 4E**). Interestingly, while patient-derived CAFs showed a similar trend of enhanced TEV uptake from CRIPTO-expressing cells relative to shCR TEV, we observed comparable levels of activated nuclear SMAD localization in CAFs treated with shCon TEVs versus those treated with shCr TEVs (**Fig 4F**). This suggests CAFs, in contrast to normal/naïve resident fibroblasts, may have activated secondary mechanisms to sustain SMAD activation independently of CRIPTO-TEV signaling (**Fig 4F**).

## DISCUSSION

In the TME, non-autonomous signaling drives adaptive responses in both stromal and parenchymal cells, contributing to cancer progression and resistance to treatment^1,12^. Our study uncovers a novel instructive role for tumor cell-derived TEVs bearing CRIPTO protein in this dynamic activity. CRIPTO-laden TEVs program fibroblasts toward an activated fibroblast cell-state, which in turn enhances tumor cell invasion. This process involves NODAL-dependent activation of SMAD2/3 in recipient fibroblasts and CRIPTO-dependent but NODAL-independent TEV uptake.

CRIPTO signaling is potentiated by direct interaction with cell surface GRP78 ^28^. GRP78 predominantly localizes to the endoplasmic reticulum where it is involved in the unfolded protein response, however, our prior studies point to stress-dependent activation of CRIPTO signaling in parenchymal cells enhancing tumor cell fitness as a function of GRP78 cell surface localization within stressed tumor microdomains^9^. We show here that such domains also exhibit colocalization of CRIPTO with EV marker CD81 and correlated increases in elongated, αSMA-positive fibroblasts and fibrosis. Chronic exposure of fibroblasts to CRIPTO-bearing EVs drives SMAD activation, the expression of classical CAF markers, and transcriptional changes associated with the TGF-β pathway, matrisome remodeling, and ER stress.

ER stress pathways have been proposed as critical for fibroblast reprogramming to various CAF-like cell states. The PERK-eIF2α-ERK1/2 axis promotes pancreatic CAF cell-state switches and ATF4 depletion in fibroblasts reduces CAF activation and fibrotic remodeling in melanoma and pancreatic murine models^29,30^. Analogous CAF pathways have yet to be described in breast cancer, however, our work suggests CRIPTO may signal in dichotomous but coordinated ways to promote or inhibit cell differentiation in separate cell compartments by responding to and modulating stress signaling.

Whether CRIPTO’s role in autocrine/paracrine signaling via EVs involves the packaging of other pathway components as EV cargo and whether these reflect the activity of previously reported interactors remains to be determined. EVs are known to promote αSMA expression in fibroblasts by transferring latent TGF-β complexes from tumor cells^19^. Furthermore, CRIPTO not only facilitates NODAL ALK4 complex formation, but also regulates proper NODAL localization and trafficking via coupling with convertases FURIN and PACE4^31,32^. In addition to TGF-β pathways, CRIPTO influences growth factor signaling through c-Src and ERK/AKT pathways^2^. The promotion of ERK activation in target fibroblast cells by non-EV-specific soluble factors that is blocked by CRIPTO-EVs suggests complex cooperation between cell types and signaling pathways. The restoration of ERK activation by inhibiting ALK4,5,7 indicates EV-based CRIPTO may balance differentiation and proliferation cues to facilitate CAF activation.

Our initial indication of CRIPTO’s role in modulating fibroblast behavior stemmed from the correlation between a CAF signature and the prevalence of CRIPTO-regulated tumor populations. We interpret this to reflect tumor sub-populations acting as key players in the complex interplay of signals between CAFs and tumor cells, highlighting the importance of understanding tumor cell heterogeneity and the nuanced “call and response” mechanisms driving tumor progression. We also noted the correlation of CRIPTO populations with a CD4^+^ T cell signature. The basis of this relationship will require further investigation, but we note that CAFs have been reported to alter intratumoral immune cell levels through cytokine secretion and alteration of the mechanical profile of the extracellular matrix^33,34^. Whether CRIPTO activity influences immune components of the TME directly or through altered CAF activity presents intriguing avenues for future research, especially regarding CRIPTO inhibition’s potential synergy with immune checkpoint blockade.

Our study utilizes various cell lines and patient-derived TNBC models highlighting elevated CRIPTO activity in this subtype. However, the findings are potentially relevant to other breast cancer subtypes and cancers in other tissues with significant fibroblast involvement, such as lung, liver, prostate, and pancreatic cancer. Indeed, CRIPTO overexpression has been reported in a broad spectrum of breast cancers, including hormone receptor (HR+) and HER2+ tumors in addition to TNBC^7^. Further, CRIPTO/GRP78 signaling has now been implicated in an array of tumor types outside of the mammary system^35,36^.

TEVs from MDA-MB-231 sublines prime pre-metastatic niches, specifically targeting cells, including S100A4+ lung fibroblasts through differential integrin expression^15^. Here we report for the first time that modulation of CRIPTO expression in tumor cells significantly changes TEV uptake rates and cellular programming of recipient cells both *in vitro* and *in vivo*. This discovery may open new avenues for therapy development, either by blocking critical CRIPTO-EV’s role in modifying stromal fibroblasts or leveraging the exceptional uptake of CRIPTO-laden EVs for stromal delivery of anticancer agents. Previous efforts to target the tumor microenvironment (TME) and CAFs in breast cancer, particularly using TGF-β inhibitors, have failed early clinical trials due to significant off-target toxicities^37^. In contrast, CRIPTO, minimally expressed in normal adult tissue, could present a safer target, with previous *in vivo* targeting showing no adverse systemic effects^9,38^. This could be especially valuable in the TNBC subtype that currently has limited targeted therapies.

In conclusion, our study establishes CRIPTO as a pivotal component of heterotypic tumor:fibroblast signaling within TNBC, with potential biologic and therapeutic relevance to other cancers via its role in modulating EV internalization and signaling within the TME. While significant advances have been made in targeting the immune stromal component of the microenvironment, limited inroads have been made in targeting CAFs in breast cancer despite their prevalence and critical roles. By targeting tumor-supportive CAFs, it may be possible to dampen mechanisms of resistance, and therefore, CRIPTO-directed therapies may prove to be important players in combinatorial therapeutic approaches. Collectively, these data have implications for biomarker and therapeutic target discovery, and the development of therapies that modulate EV function via CRIPTO.

## METHODS

### Cell culture

Authenticated triple-negative breast cancer cells, MDA-MB-468 (RRID: CVCL_0419), and MDA-MB-231 (RRID: CVCL_0062), and murine NIH-3T3 (RRID: CVCL_0594) cells were acquired from ATCC. Cells were cultured in ‘Tumor Media’ (Dulbecco’s modified Eagle’s media (DMEM), 10% fetal bovine serum, and ciprofloxacin 10μg/mL). NIH-3T3 cells were treated with 1μg/mL of mitomycin-C for 2 hours to generate growth arrested fibroblasts. For treatments with inhibitors or EVs, cells were cultured with media containing EV-depleted FBS for 24 hours prior to treatment. PDxOs were a generous gift from the Welm Labs (HCI) and were cultured according to their published protocol^39^. Primary human mammary fibroblasts and CAFs were aquired from Cell Biologics and cultured in tumor media tumor media supplemented with FGF (100ng/mL) and hydrocortisone (1μg/mL).

### RNA-sequencing

For scRNA-seq MDA-MB-468 expressing either s14-Con or s14-A4 were grown as organoids in low adhesion plates with 5% Engelbreth-Holm-Swarm (EHS)-derived basement membrane matrix for 7 days. Organoids were trypsinized, dissociated with dispase and barcoded with the Chromium Next GEM Single Cell 3’ Kit v3.1 with cell multiplexing according to the manufacturer’s instructions [Protocol #1] (#100261, 10X Genomics, Pleasanton, CA). Sequencing was performed on the NovaSeq 6000 with 150×150 paired end mode by the University of Utah High Throughput Genomics Core (HTGC) at Huntsman Cancer Institute based on the 10X genomics user guide (CG000315).

For bulk RNAsequencing, NIH/3T3s were cultured in EV-depleted tumor media and treated with 2.5μg of purified EVs from MDA-MB-468 shCon, shCripto, or CRIPTO overexpressing cells for 24 hours. RNA was collected with RNeasy Kit according the manufacturer’s instructions (#74104, Qiagen, Germantown, MD) and purity was validated using a 2100 Bioanylyzer (Agilent Technologies, Santa Clara, CA) and Qubit^TM^ Broad Range RNA kit (Thermo Fisher Scientific, Waltham, MA). Libraries were prepped with the NEBNext Ultra II Directional RNA library Prep with Poly(A) isolation (New England Biolabs, Ipswich, MA). Reads were aligned to the mouse genome using Elysium. We quantified gene expression as transcripts per million (TPM).

### Lentiviral Vectors

S14-Con and -A4 lentiviral expression vectors are previously described tet-inducible constructs^9^. *TDGF1* and *NODAL* knockdown constructs were purchased from Horizon Discovery (cat #V3SH11255-01EG6997 Cambridge, UK) and Origene (cat #TL501492 Rockville, MD) respectively.

### Extracellular Vesicle Isolation and labeling

EVs were isolated by differential ultracentrifugation as previously described and validated according to recent MISEV2023 guidlines^23,40^. Cells were cultured at 70-90% confluency for 48 hours in Tumor Media with EV-depleted FBS. Conditioned media was collected and subjected to sequential centrifugation at 500g for 10 minutes, 2500g for 20 minutes, 10000g for 30 minutes, passed through 0.2μm PES filters and centrifuged at 120,000g for 2 hours (SW28 rotor, K=246). Pelleted EVs were resuspended in 0.2 μm filtered PBS and centrifuged at 120,000g for 2 hours (SW55Ti rotor, K=48). The resulting EV pellet was resuspended in either filtered PBS or RIPA depending on downstream assays. For PKH26 staining, isolated EVs were labeled according to the manufacturer’s instructions (MIDI6-1KT, Sigma, Burlington, MA). Excess PKH was removed by centrifugation at 190,000g for 2 hours with a 0.971M sucrose cushion (SW55Ti rotor, K=48). EVs were resuspended in 9mL of PBS and filtered with a 10kDa centrifugal filter (UFC901008, Sigma, Burlington, MA) at 3000g for 30 minutes. PKH26 labeled EVs were stored at 4C and used within 24 hours of labeling.

### Nanoparticle Tracking Analysis

EV samples were diluted 1:20 with 0.2μm filtered PBS. Size distribution and concentration were analyzed on a NS300 (Malvern, Malvern UK). Samples were recorded blinded over 3 videos of 60s at detection threshold: 3, camera level: 13 with a coefficient of variance of <15%.

### Electron Microscopy

Cryogenic electron microscopy was performed by the electron microscopy core at the University of Utah. Briefly, EVs were applied (3.5 μL) to glow-discharged 2/1 Quantifoil carbon 300-mesh Cu grids. Grids were blotted for 4-6 seconds on the front-side of the grid and immediately plunged into liquid Ethane using a Leica plunge freezing device.

### EV Uptake Timecourse

Cells were grown in 8 well chambered coverglass (#155409, Thermo Fisher Scientific, Waltham, MA) and imaged with an SP8 white laser confocal (Leica Biosystems, Wetzlar, Germany) with a stage top incubator (Okolab, Sewickley, PA). Two images per well were taken at 200X magnification in 20 minutes intervals over an 8 hour period. 1μg of PKH26-labeled EVs were added after an initial image and fluorescence intensity was normalized based on the fluorescent EV signal after the secod timepoint for each condition.

### EV delivery of CRE

EVs were isolated from from control and CRIPTO overexpressing 293A cells, and for each treatment, 3ug of purified EVs were electroporated in 20μL of SF buffer (#V4XC-2032, Lonza, Basel, Switzerland) with 0.4μg of CRE-expressing plasmid (Addgene #172433) on a 4D-Nucleofector (Lonza, Basel, Switzerland) using program EO-100. Electroporated EVs were added to 293T switch reporter cells carrying a floxed red-to-green fluorescent protein reporter in 200μL of media, and were incubated for at least 72 hours prior to imaging.

### In Vivo Studies

10-12 week old SCID-Beige mice were purchased from Charles River Laboratory (Wilmington, MA). All animal care and procedures were approved and monitored by an Institutional Animal Care and Use Committee. For EV uptake studies, mice received retroorbital injections of 10μg of PKH26 (MIDI6-1KT, Sigma, Burlington, MA) labeled EV as previously described^15^. 24 hours following EV administration, animals were euthanized, and lungs and livers were harvested, fixed in 4% PFA overnight followed by sequential incubation in 10% sucrose (1 hour), 20% sucrose (1 hour), and 30% sucrose (overnight) solutions. Fixed tissues were embedded in O.C.T, frozen, and cryosectioned at 5μm.

### Immunofluorescence

Cells were cultured on 8-well chamber slides with removable covers (#155409PK, Thermo Fisher Scientific, Waltham, MA). Upon culture endpoint cells were fixed with 4% PFA, permeabilized with 0.05% Triton-X 100, and blocked with 10% goat serum. For EV internalization, cells were incubated with 1μg of PKH26-labeled EVs for 1 hour prior to fixation. Permeabilized cells were probed for SMAD2/3 (#AF3797, R&D Biosystems, 10μg/mL) and imaged on an SP8 white laser confocal microscope (Leica Biosystems, Wetzlar, Germany). The following antibodies were used for additional immunofluorescence assays: CRIPTO (ab19917, Abcam, 1:800), CD81 (Clone 1.3.3.22, Thermo Fisher Scientific, Waltham, MA, 1:20), Goat anti-mouse AF647 (ab150115, Abcam, 1:300), Goat anti-rabbit 568 (A-11011, Thermo Fisher Scientific, Waltham, MA, 1:200).

### Microwell Organoid Assay

MDA-MB-468 and mitomycin-C treated NIH-3T3 cells were plated at a ratio of 10:1 in AggreWell^TM^400 plates at approximately 50 cells/microwell according to the manufacturer’s instructions (#34415, Stemcell Technologies, Vancouver, BC) and grown for 5 days in tumor media containing 1μg/mL doxycycline. Dispersed cell counts were measured blinded as the number of cells present beyond the average diameter of MDA-MB-468 monoculture spheroids using the image analysis software FIJI^41^.

### Western Blots

Cells and EVs were lysed in RIPA buffer supplemented with protease/phosphatase inhibitors (#78440, Thermo Fisher Scientific, Waltham, MA) and quantified with DC Protein Assay (#500111, BioRad, Hercules, CA). Lysates were boiled in LDS loading buffer with 0.1M DTT and run on 4-12% precast Bis-Tris gels (NP0336BOX, Thermo Fisher Scientific, Waltham, MA). Proteins were transferred using an iBlot2 transfer device (Thermo Fisher Scientific, Waltham, MA), blocked with Intercept Blocking Buffer (927-7001, LiCor Biosciences, Lincoln, NE), and immunoblotted with the following antibodies: phospho-SMAD2 (Ser465/467)/SMAD3 (Ser423/425) (Clone D27F4, Cell Signaling, 1:1000), SMAD2/3 (#AF3797, R&D Biosystems, 0.5μg/mL), Phospho-ERK1/2 (Thr202/204) (Clone D13.144E, Cell Signaling, 1:1000), ERK1/2 (L34F12, Cell Signaling, 1:2000), Vinculin (Clone EPR8185, Abcam, 1:1000), CRIPTO (ab19917, Abcam, 1:800; FigS1F CRIPTO-1 Mouse Monoclonal from David Salomon), Calnexin (Clone C5C9, Cell Signaling, 1:1000), HSP70 (#4872, Cell Signaling, 1:1000), Flotillin-1 (Clone D2V7J, Cell Signaling, 1:1000), CD81 (Clone D3N2D, Cell Signaling, 1:1000), NODAL (Clone WS65, Santa Cruz Biotechnology, 1:200), Goat anti-rabbit IgG AF680 (#A21076, Thermo Fisher Scientific, 1:2000), Donkey anti-goat IgG AF800 (#A32930, Thermo Fisher Scientific, 1:2000), Goat anti-mouse IgG (#A11375, Thermo Fisher Scientific, 1:2000).

### PCR

RNA was isolated from cells using the RNeasy Kit (#74104, Qiagen, Germantown, MD) and quantified on a NanoDrop 8000 (Thermo Fisher Scientific, Waltham, MA). RT-qPCR was performed using PowerUp^TM^ Sybr^TM^ Green Master Mix (#A25741, Thermo Fisher Scientific, Waltham, MA) according to manufacturer’s protocol and utilized a BioRad^TM^ CFX384 thermocycler. Comparisons were made using the following pre-validated housekeeping genes: *POLR2A* (Hs.PT.39a.19639531; IDT, San Diego, CA); *Polr2a* (MM.PT.39a.22214849; IDT, San Diego, CA). The following custom RNA-specific primer sequences were designed: *TDGF1:* (Forward: TCCTTTTGTGCCTGCCCTC Reverse: CACAGGGAACACTTCTTGG); *SOSTDC1* (Forward: TCAAGCCAGAAATGGAGGCAGG Reverse: GCCATCAGAGATGTATTTGGTGG); *CRYAB* (Forward: ACTTCCCTGAGTCCCTTCTACC Reverse: GGAGAAGTGCTTCACATCCAGG); *Tnc* (Forward: GAGACCTGACACGGAGTATGAG Reverse: CTCCAAGGTGATGCTGTTGTCTG); *TGF-β* (Forward: TGATACGCCTGAGTGGCTGTCT Reverse: CACAAGAGCAGTGAGCGCTGAA); *S100a4* (Forward: AGCTCAAGGAGCTACTGACCAG Reverse: GCTGTCCAAGTTGCTCATCACC); *Pdgfr*α (Forward: GCAGTTGCCTTACGACTCCAGA Reverse: GGTTTGAGCATCTTCACAGCCAC); *Fn1* (Forward: CCCTATCTCTGATACCGTTGTCC Reverse: TGCCGCAACTACTGTGATTCGG); *Fap* (Forward: CCCGCGTAACACAGGATTCACT Reverse: CACACTTCTTGCTCGGAGGAGA); *Acta2* (Forward: CACTATTGGCAACGAGCGC Reverse: CCAATGAAGGAAGGCTGGAA).

### Invasion Assays

Invasion assays were performed using 6.5mm polycarbonate membrane Transwell^®^ inserts with 8.0μm pores. Inserts were coated with 45μL of EHS basement membrane matrix and solidified at 37C for 5 minutes^42^. 2.0×10^4^ tumor cells plated in 100 μL on top of the EHS coated insert were grown with fibroblasts for 72 hours to prime fibroblasts. Priming inserts were discarded and fresh EHS coated inserts with 2.0×10^4^ tumor cells were placed on top of both naïve and primed fibroblasts and cultured for an additional 48 and 72 hours for cell lines and PDxOs, respectively. Inserts were washed with PBS, fixed with 4% PFA for 15 minutes at RT, permeabilized with 0.05% Triton X-100, incubated with DAPI 1 μg/mL for 5 minutes. EHS matrix/non-migrated cells were removed from the top of the transwell with a cotton tipped applicator following permeabilization. Membranes were excised, mounted, and coverslipped in ProLong^TM^ Gold mounting media (#P36930, Thermo Fisher Scientific, Waltham, MA), and imaged on an Axioscan 7 Slide Scanner (Zeiss, Oberkochen, Germany). Membrane area was manually measured and DAPI counts were performed using the automated cell counting program in QuPath^43^.

## DATA AVAILABILITY STATEMENT

The data generated in this study are available in Gene Expression Omnibus (GEO) (GSE262955; GSE262956), or upon reasonable request from the corresponding author.

## ACKNOWLEDGMENTS

We would like to thank the cell imaging, electron microscopy, and high throughput genomics cores at HCI for their expertise and equipment, and R. O’Connell for generous use of the nanosight.

## AUTHOR CONTRIBUTIONS

Concept and writing: DWF and BTS wrote the manuscript with input from all the authors. DWF and BTS conceived and designed the study, and DWF and BLG conducted the experiments. HEG and MDS performed blinded analysis of PDX sections and microwell assays, respectively.

