## Supplemental Table 1 for "CRIPTO promotes extracellular vesicle uptake and activation of cancer associated fibroblasts"

**Table S1. Differentially expressed genes from cluster 5 and 8**

| Gene | Fold Change | Cluster | Signature Weight |
|---|---|---|---|
| AL583785.1 | 4.92 | 8 | 4.92 |
| AARD | 1.93 | 8 | 1.93 |
| IGF2BP3 | 3.30 | 8 | 3.30 |
| TFCP2L1 | 2.98 | 8 | 2.98 |
| CRYAB[#] | 1.63 | 8 | 1.63 |
| TNS3 | 2.49 | 8 | 2.49 |
| SAA1[#] | 2.23 | 8 | 2.23 |
| TCF7L1 | 2.04 | 8 | 2.04 |
| SYNJ2 | 1.97 | 8 | 1.97 |
| PEG10 | 1.97 | 8 | 1.97 |
| LINC01619 | 1.97 | 8 | 1.97 |
| AUTS2 | 1.93 | 8 | 1.93 |
| SMYD3 | 1.89 | 8 | 1.89 |
| SORBS1 | 1.85 | 8 | 1.85 |
| OSBP2 | 1.84 | 8 | 1.84 |
| KIF26B | 1.72 | 8 | 1.72 |
| CRIM1[#] | 1.57 | 8 | 1.57 |
| LDLRAD4 | 1.54 | 8 | 1.54 |
| TMPRSS2[#] | 1.51 | 8 | 1.51 |
| HLA-B[#] | 1.49 | 8 | 1.49 |
| GSTM3[#] | 1.47 | 8 | 1.47 |
| ATP6V0A4[#] | 1.41 | 8 | 1.41 |
| RERG | 1.39 | 8 | 1.39 |
| SYT12 | 1.37 | 8 | 1.37 |
| EPB41L2*[#] | 1.36 | 8 | 1.36 |
| PROM1 | 1.36 | 8 | 1.36 |
| UTRN[#] | 1.34 | 8 | 1.34 |
| CHODL | 1.34 | 8 | 1.34 |
| AC008014.1 | 1.28 | 8 | 1.28 |
| ANKH | 1.28 | 8 | 1.28 |
| TGFB2 | 1.26 | 8 | 1.26 |
| RAI14 | 1.26 | 8 | 1.26 |
| PDGFC[#] | 1.25 | 8 | 1.25 |
| L3MBTL4 | 1.23 | 8 | 1.23 |
| FBXO32 | 1.22 | 8 | 1.22 |
| SHROOM3 | 1.21 | 8 | 1.21 |
| MAGI1 | 1.17 | 8 | 1.17 |
| BTG1* | 1.14 | 8 | 1.14 |
| ST8SIA1 | 1.14 | 8 | 1.14 |
| LGR4 | 1.13 | 8 | 1.13 |
| MBOAT1 | 1.11 | 8 | 1.11 |
| IGFL2-AS1 | 1.11 | 8 | 1.11 |

| | | | |
|---|---|---|---|
| GAB2 | 1.10 | 8 | 1.10 |
| EFHD1[#] | 1.09 | 8 | 1.09 |
| PC | 1.09 | 8 | 1.09 |
| ANXA1[#] | -1.43 | 8 | -1.43 |
| OSBPL6 | -1.56 | 8 | -1.56 |
| LGALS1[#] | -1.65 | 8 | -1.65 |
| SERPINA3[#] | -1.69 | 8 | -1.69 |
| PLXDC2[#] | -1.74 | 8 | -1.74 |
| LINC00607 | -1.75 | 8 | -1.75 |
| OBP2B | -1.84 | 8 | -1.84 |
| CAMK2N1 | -1.87 | 8 | -1.87 |
| ID1 | -1.91 | 8 | -1.91 |
| CA9 | -2.02 | 8 | -2.02 |
| CLDN10-AS1 | -2.15 | 8 | -2.15 |
| AREG[#] | -2.31 | 8 | -2.31 |
| KCNMA1^[#] | -2.48 | 8 | -2.48 |
| UBE2C* | -4.10 | 5 | 4.10 |
| TOP2A^ | -3.58 | 5 | 3.58 |
| DEPDC1^ | -3.15 | 5 | 3.15 |
| NDC80* | -3.31 | 5 | 3.31 |
| GTSE1^ | -3.21 | 5 | 3.21 |
| KIF2C*^ | -3.00 | 5 | 3.00 |
| CDK1* | -3.47 | 5 | 3.47 |
| ASPM | -2.42 | 5 | 2.42 |
| CENPF*^ | -1.92 | 5 | 1.92 |
| DLGAP5 | -2.33 | 5 | 2.33 |
| AURKA*^ | -2.55 | 5 | 2.55 |
| HIST1H1B | -2.87 | 5 | 2.87 |
| TPX2*^ | -2.57 | 5 | 2.57 |
| HIST1H1D | -0.80 | 5 | 0.80 |
| MKI67*^ | -2.34 | 5 | 2.34 |
| HMMR^ | -2.48 | 5 | 2.48 |
| HIST1H4C | -2.70 | 5 | 2.70 |
| KIF4A^ | -2.05 | 5 | 2.05 |
| CDC20*^ | -2.52 | 5 | 2.52 |
| CKS1B* | -2.38 | 5 | 2.38 |
| NUSAP1^ | -2.51 | 5 | 2.51 |
| KIF20B* | -1.77 | 5 | 1.77 |
| HMGB2 | -2.20 | 5 | 2.20 |
| CKS2 | -1.71 | 5 | 1.71 |
| DIAPH3 | -2.70 | 5 | 2.70 |
| ANLN^ | -1.92 | 5 | 1.92 |
| CCNB1* | -1.82 | 5 | 1.82 |
| PRC1*^ | -2.19 | 5 | 2.19 |

| | | | |
|---|---|---|---|
| CCNB2*^ | -0.31 | 5 | 0.31 |
| BIRC5*^ | -1.80 | 5 | 1.80 |
| HIST1H1A | -1.73 | 5 | 1.73 |
| HIST2H2AC | -0.67 | 5 | 0.67 |
| RRM2^ | -2.74 | 5 | 2.74 |
| UBE2S* | -1.97 | 5 | 1.97 |
| CENPA^ | -0.80 | 5 | 0.80 |
| KPNA2 | -0.32 | 5 | 0.32 |

*Proliferation genes
^Shared genes with CAF signature
#Enriched in Exosomes
