## Supplementary Figures for "CRIPTO promotes extracellular vesicle uptake and activation of cancer associated fibroblasts"

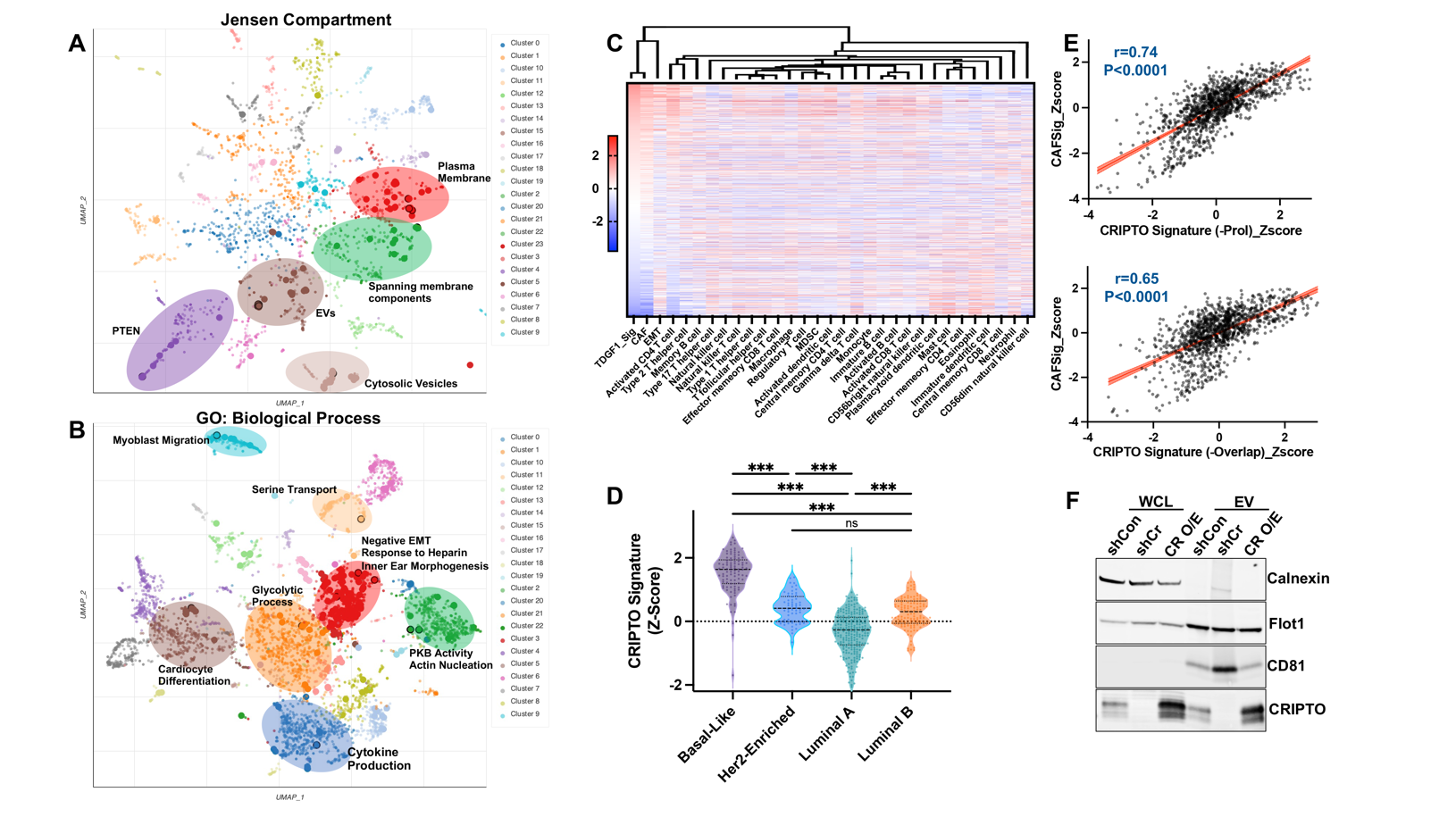
**Figure S1. CRIPTO is associated with TEV signaling and breast CAF levels**

UMAPS of (A) Jensen compartment terms used in Fig 1D and (B) GO biological processes, outlined points denote significant groups (q<0.1; Benjamini-Hochberg adjusted). (C) Hierarchical clustering of relative CRIPTO signature and published gene signatures expression from Fig 1E across breast cancer TCGA data. (D) CRIPTO signature enrichment across PAM50 subtypes^45^, one-way ANOVA with Tukey’s multiple comparisons (E) Pearson correlation of the CAF and CRIPTO signatures after removing genes annotated as ‘proliferation’ or ‘cell division’ by GO (top) and shared between the signatures (bottom). (F) Western blot of whole cell lysates (WCL) and EVs from MDA-MB-468 shCon, shCr, and Cr O/E cells.

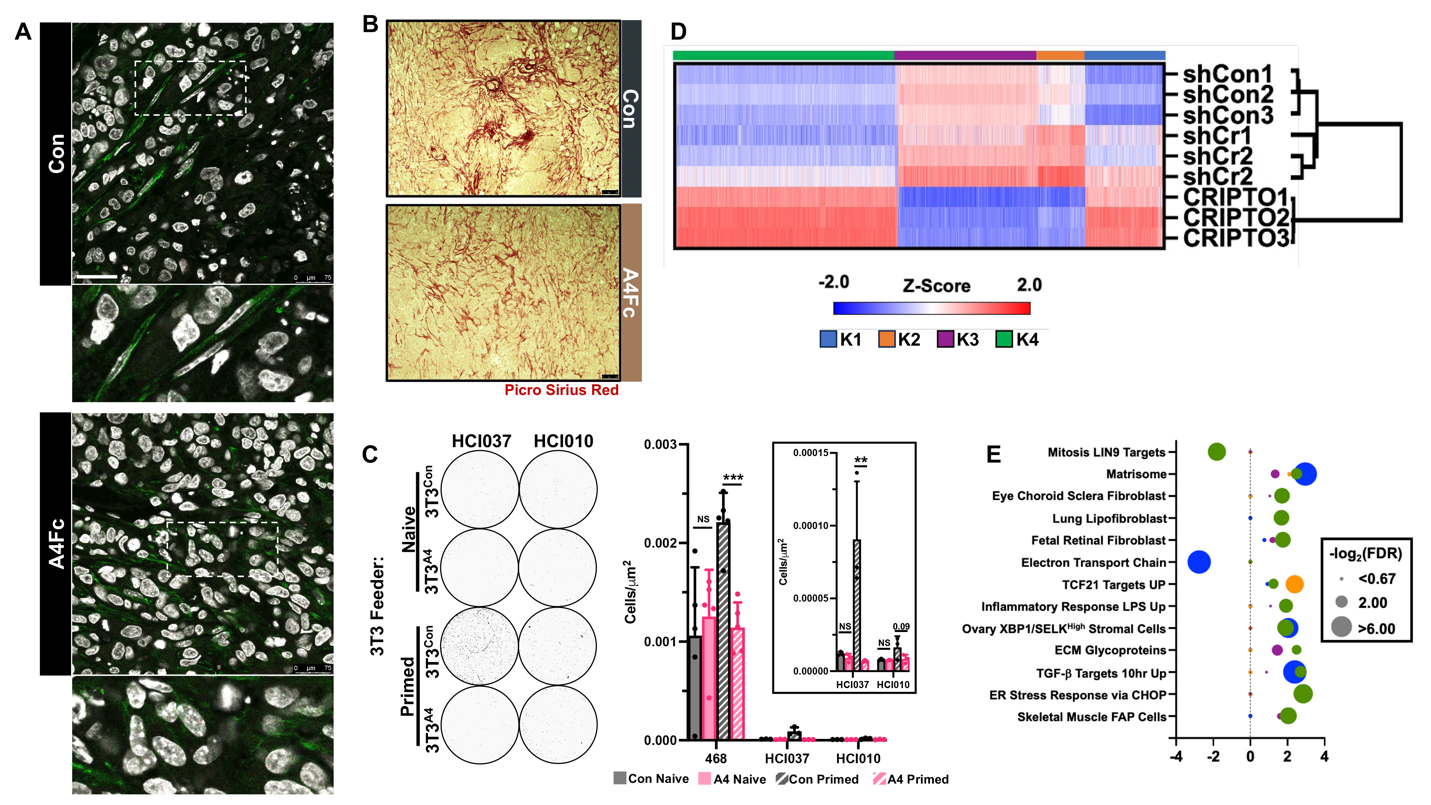
**Figure S2. Paracrine CRIPTO signaling promotes fibroblast programming and cell-state switches.**

MDA-MB-468 s14-con and s14-A4Fc expressing xenografts stained for (A) αSMA (scale bar=75μm) and (B) collagen expression (scale bar=100μm). (C) Representative PDxO invasion assays and unadjusted invasion levels from Fig 3D. (D) Hierarchical and k-means clustering of bulk RNAseq of 3T3 cells treated with TEVs from MDA-MB-468 shCon, shCr, or Cr O/E cells. (E) Significant GSEA sets for each cluster identified in panel D; colors correspond to cluster colors.
